# Predicting age from cortical structure across the lifespan

**DOI:** 10.1101/248518

**Authors:** Christopher R. Madan, Elizabeth A. Kensinger

**Affiliations:** School of Psychology, University of Nottingham, Nottingham, UK; Department of Psychology, Boston College, Chestnut Hill, MA, USA

**Keywords:** aging, structural MRI, brain morphology, fractal dimensionality, cortical complexity, gyrification

## Abstract

Despite inter-individual differences in cortical structure, cross-sectional and longitudinal studies have demonstrated a large degree of population-level consistency in age-related differences in brain morphology. The present study assessed how accurately an individual’s age could be predicted by estimates of cortical morphology, comparing a variety of structural measures, including thickness, gyrification, and fractal dimensionality. Structural measures were calculated across up to seven different parcellation approaches, ranging from 1 region to 1000 regions. The age-prediction framework was trained using morphological measures obtained from T1-weighted MRI volumes collected from multiple sites, yielding a training dataset of 1056 healthy adults, aged 18-97. Age predictions were calculated using a machine-learning approach that incorporated non-linear differences over the lifespan. In two independent, held-out test samples, age predictions had a median error of 6-7 years. Age predictions were best when using a combination of cortical metrics, both thickness and fractal dimensionality. Overall, the results reveal that age-related differences in brain structure are systematic enough to enable reliable age prediction based on metrics of cortical morphology.

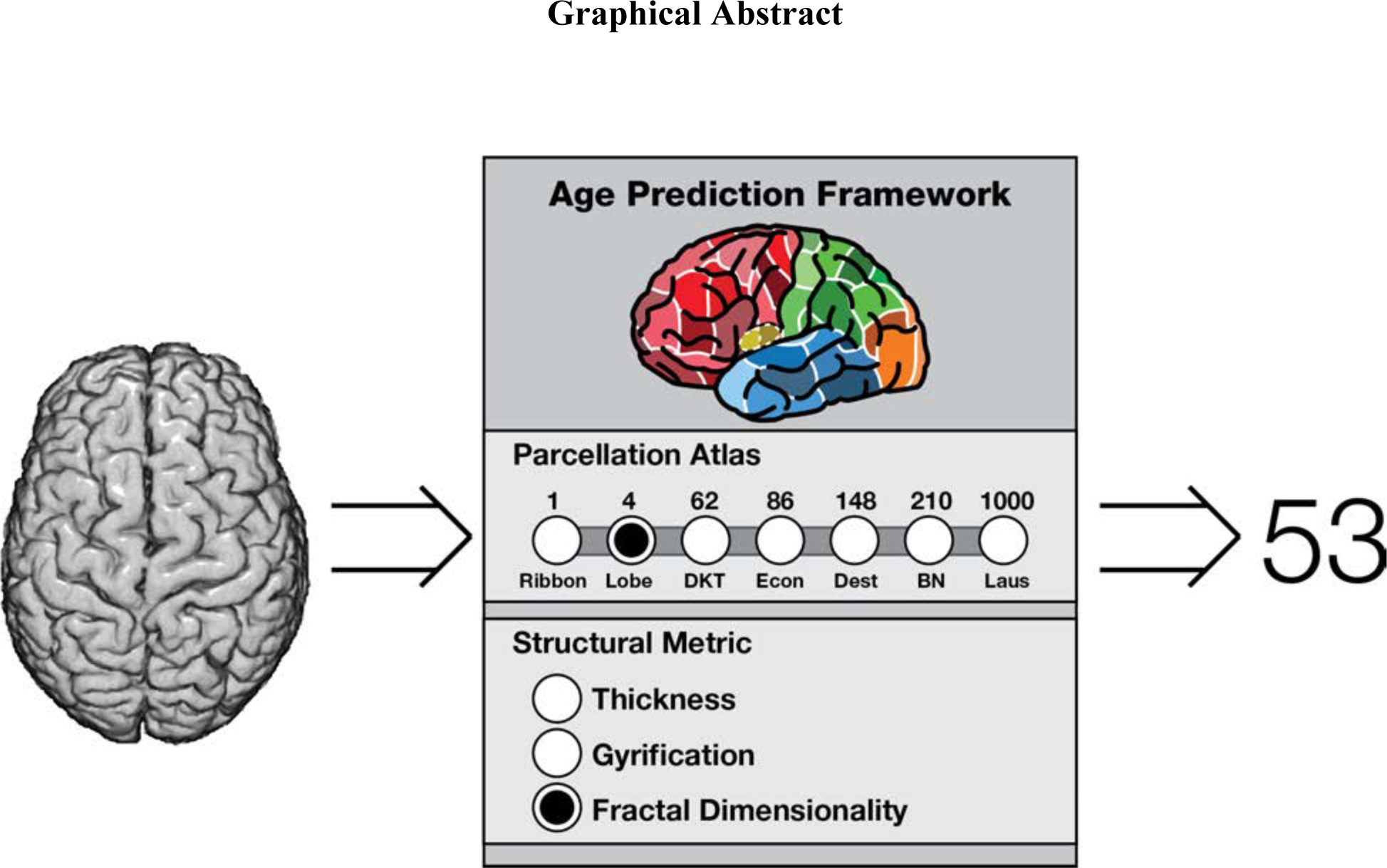
Graphical Abstract.

Several measures of cortical structure differ in relation to age. We examined the cortical granularity of these differences across seven parcellation approaches, from a 1 region (unparcellated cortical ribbon) to 1000 regions (patches with boundaries informed by anatomical landmarks), and three measures: thickness, gyrification, and fractal dimensionality. Rather than merely examining age-related relationships, we examined how these parcellations and measures can be used to *predict* age.

## Introduction

It is well-established that the structure of the brain changes as adults age—with decreases in cortical thickness as one of the most pronounced of these changes (e.g., Fjell et al., 2009, 2013; Hogstrom et al., 2013; Hutton et al., 2009; Irimia et al., 2015; Jernigan et al., 2001; Lemaitre et al., 2012; Madan & Kensinger, 2016; McKay et al., 2014; Raz & Rodrigue, 2006; Salat et al., 2004). However, other measures of cortical structure are also sensitive to age-related differences, such as gyrification (Cao et al., 2017; Hogstrom et al., 2013; Jockwitz et al., 2017; Madan & Kensinger, 2016; Magnotta et al., 1999; Wang et al., 2016), which is a ratio of the regional surface area relative to the surface area of a simulated enclosing surface (Armstrong et al., 1995; Hofman, 1991; Kochunov et al., 2012; Toro et al., 2008; Zilles et al., 1988, 1989). More recently, a mathematical measure of the complexity of a structure, fractal dimensionality, has also been shown to index age-related differences in brain structure (Madan & Kensinger, 2016). While the use of fractal dimensionality with cortical aging is recent, it has been used in prior studies investigating differences in cortical structure in patient populations (Cook et al., 1995; Free et al., 1996; King et al., 2009, 2010; Nenadic et al., 2014; Sandu et al., 2008; Thompson et al., 2005; Wu et al., 2010), as well as cross-species comparisons (Hofman, 1991). Importantly, age-related differences in these structural measures are not homogenous across the cortex; decreases in cortical thickness are most evident in frontal regions, while gyrification decreases primarily in parietal cortex. Given this, we wondered what degree of precision is useful in understanding age-related differences in cortical structure. Furthermore, it is unknown how well the relation between age and cortical structure metrics will generalize across independent samples. In the present study we sought to examine (1) the relative sensitivity of different cortical measures to age-related differences across the adult lifespan, (2) the granularity of these differences across different cortical parcellation approaches, and (3) how well these different measures and parcellations can be used to *predict* age in independent samples. These findings should further our understanding of the neurobiological basis of healthy aging (Falk et al., 2013; Reagh & Yassa, 2017).

### Cortical metrics and granularity

While there is heterogeneity in age-related differences in cortical structure—e.g., greater cortical thinning in frontal cortex than occipital (Allen et al., 2005; Fjell et al., 2009; Hogstrom et al., 2013; Hutton et al., 2009; Salat et al., 2004; Sowell et al., 2003)—it is unclear what degree of parcellation would be beneficial in characterizing healthy aging. Lobe-wise estimates of cortical thickness would likely be beneficial relative to overall mean cortical thickness, but would estimates for distinct gyri and lobules provide additional information? At some level of parcellation, additional predictive features should diminish, as cortical thickness between adjacent patches of cortex would be highly similar (within an individual).

Here we investigated three measures of cortical structure—thickness, gyrification, and fractal dimensionality. These measures were selected based on prior studies that had identified relationships between these measures and healthy aging, although these previous studies had used correlations rather than predictive models (e.g., Fjell et al., 2009; Hogstrom et al., 2013; Madan & Kensinger, 2016; McKay et al., 2014; Salat et al., 2004). This approach of using surface-based morphology allowed us to examine *distinct* measures of cortical structure. In particular, this is in contrast to voxel-based morphology (VBM) techniques which estimate gray matter volume and are influenced by a combination of structural features (Fairchild et al., 2015; Gerrits et al., 2016; Hutton et al., 2009; Palaniyappan & Liddle, 2012). This point is made explicit by Mechelli et al. (2005), “exactly the same differences would be detected when comparing images of thin, unfolded cortex against thin, folded cortex and thick, unfolded cortex” (see Figure 5 of Mechelli et al., 2005). As such, VBM is not sensitive to precise features as we sought to examine here, whereas surface-based morphometry captures these details unambiguously.

Many different approaches have been suggested to parcellate the human cortex (see Zilles & Amunts, 2010, for a review); here we used seven parcellation approaches, focusing on atlases that have been implemented within FreeSurfer. The two standard parcellation atlases within FreeSurfer are the Desikan-Killany-Tourville (DKT) atlas (Desikan et al., 2006; Klein & Tourville, 2012) and the Destrieux atlas (Destrieux et al., 2010), which divide the cortex into 62 and 148 parcellations, respectively. Both of these atlases define boundaries based on anatomical landmarks, with the main difference being that the Destrieux atlas divides gyri and sulci into separate parcellations, while the DKT atlas generally uses sulci as parcellation boundaries between gyri, as shown in Figure 1. As two coarse atlases, we also considered the unparcellated cortical ribbon (i.e., only 1 region) as well as a lobe-wise parcellation (4 regions). Supplementing the parcellation atlases standard within FreeSurfer, we also conducted analyses using structural metrics derived from three additional parcellation schemes: von Economo-Koskinas (86 regions; Scholtens et al., 2018), Brainnetome (210 regions; Fan et al., 2016), and Lausanne (1000 regions; Hagmann et al., 2008). The von Economo-Koskinas atlas was developed by Scholtens et al. (2018), based on the foundational work of von Economo and Koskinas (1925, 2008). This parcellation atlas is based on the cyctoarchitecture of the cortex (also see von Economo, 1927, 2009; previously some have used a hybrid approach to integrate von Economo’s work with the Desikan-Killany atlas [Scholtens et al., 2015; van den Heuvel et al., 2015]). The Brainnetome atlas takes a different approach, using the Desikan-Killany atlas and further parcellating it based on connectivity data from diffusion and resting-state scans (Fan et al., 2016). The Lausanne atlas also initially starts from the Desikan-Killany atlas and then further parcellates it into patches of approximately similar area (Hagmann et al., 2008); here we used the 1000 parcellation variant, though variants with less patches also exist. By predicting age in independent samples, across these different parcellation approaches, we can assess the cortical granularity of different structural metrics in relation to age-related differences in cortical structure.

**Figure 1.**
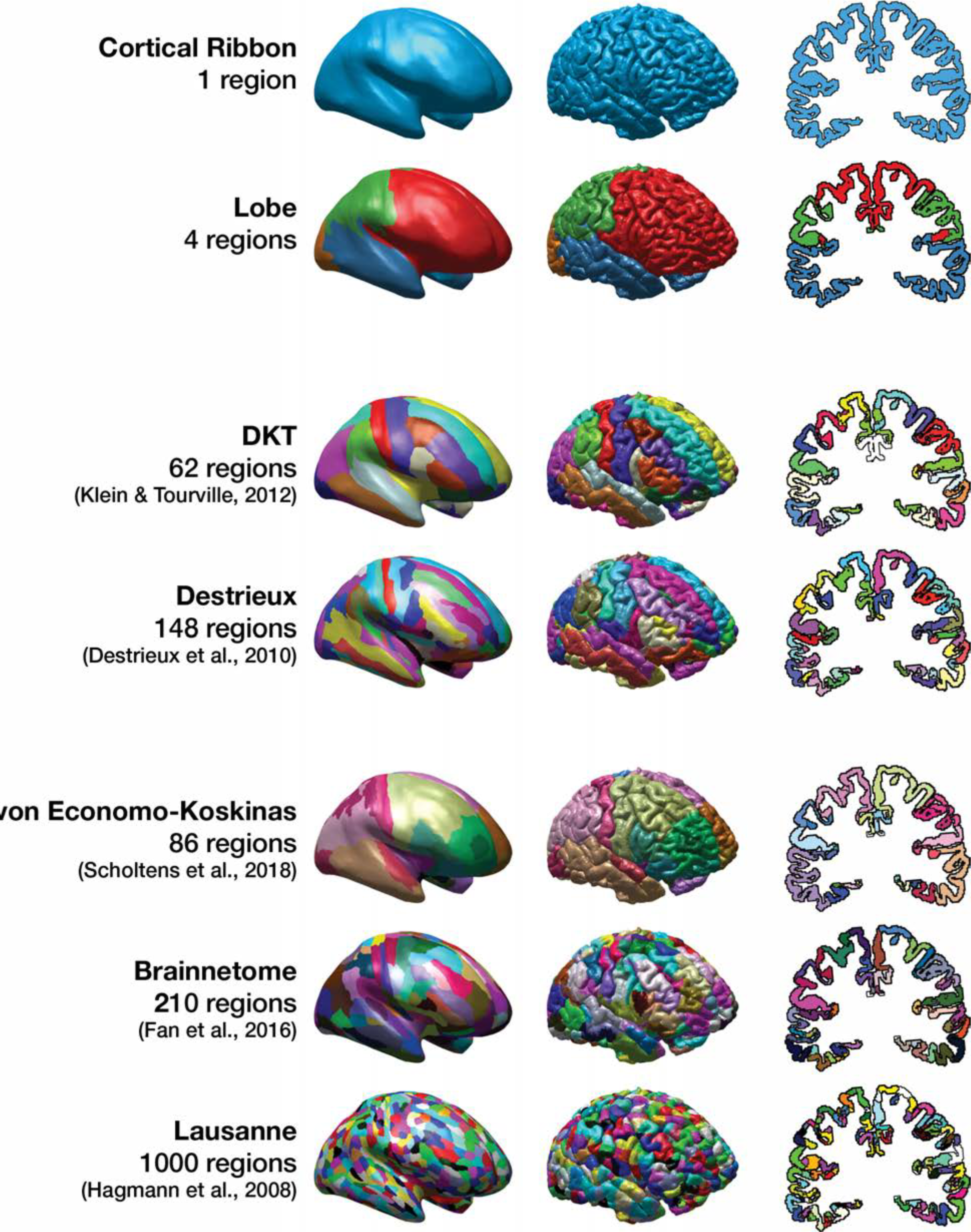
Inflated and pial surfaces and an oblique coronal slice, from a young adult (20-year-old male), illustrating the seven parcellation approaches used.

While these parcellations are defined using anatomical landmarks, different parcellation approaches exist and it is unknown to what degree more discrete cortical parcellation regions— the topological granularity—will provide additional predictive value to inform the age prediction performance. It is also unknown whether some metrics of brain morphology would yield better prediction accuracy than others; the topology of age-related differences in thickness and gyrification have been shown to differ, and thus may be indicators of distinct aging processes (Hogstrom et al., 2013; Madan & Kensinger, 2016). More recently, we have shown that measures of fractal dimensionality can show stronger age correlations than measures of cortical thickness and gyrification (Madan & Kensinger, 2016), but it was not known whether this would translate to additional predictive accuracy.

### Predicting age from cortical measures

Though age-related differences in brain morphology are robust, there also can be extensive individual variability in the trajectory of brain aging (e.g., Pfefferbaum & Sullivan, 2015), and cross-sectional comparisons demonstrate that some older adults can have similar mean cortical thickness volumes as young adults (e.g., Fjell et al., 2009; Madan & Kensinger, 2016; Salat et al., 2004). In the absence of acquiring multiple MRI scans from the same individual, there are no methods that can easily discriminate among different age-related trajectories, nor is there agreement as to which metrics might be the best for identifying which individuals are likely to be on an accelerated-aging trajectory. The present study took a first step by examining which parcellation techniques and estimates of cortical morphology would best *predict* an individual’s age. Critically, here we measured the age predictions, rather than simply correlations with age, because significant relationships are not necessarily indicative of predictive value (e.g., see Lo et al., 2015). As a further point of consideration, age-related differences in these metrics have been shown to be non-linear (Fjell et al., 2010, 2013), with age-related trajectories declining more steeply after ‘critical ages,’ that also differ across structures (generally occurring between ages 40 and 70). Based on this evidence, we used a multiple smoothing-spline based regression procedure (Madan, 2016). Figure 2 shows representative cortical surfaces for individuals with ages in the first year of each decade, for each sex and training dataset used here. These cortical surfaces visibly show the degree of inter-individual differences in cortical structure, but also make apparent the age-related differences in gyrification and sulcal width, along with other structural characteristics such as Yakovlevian torque. It is also visible here that male brains are generally slightly larger than female brains, and that brain size tends to decrease with age. (Note that these are representative individual brains from the datasets used here, however, and not ‘average’ brains.)

**Figure 2.**
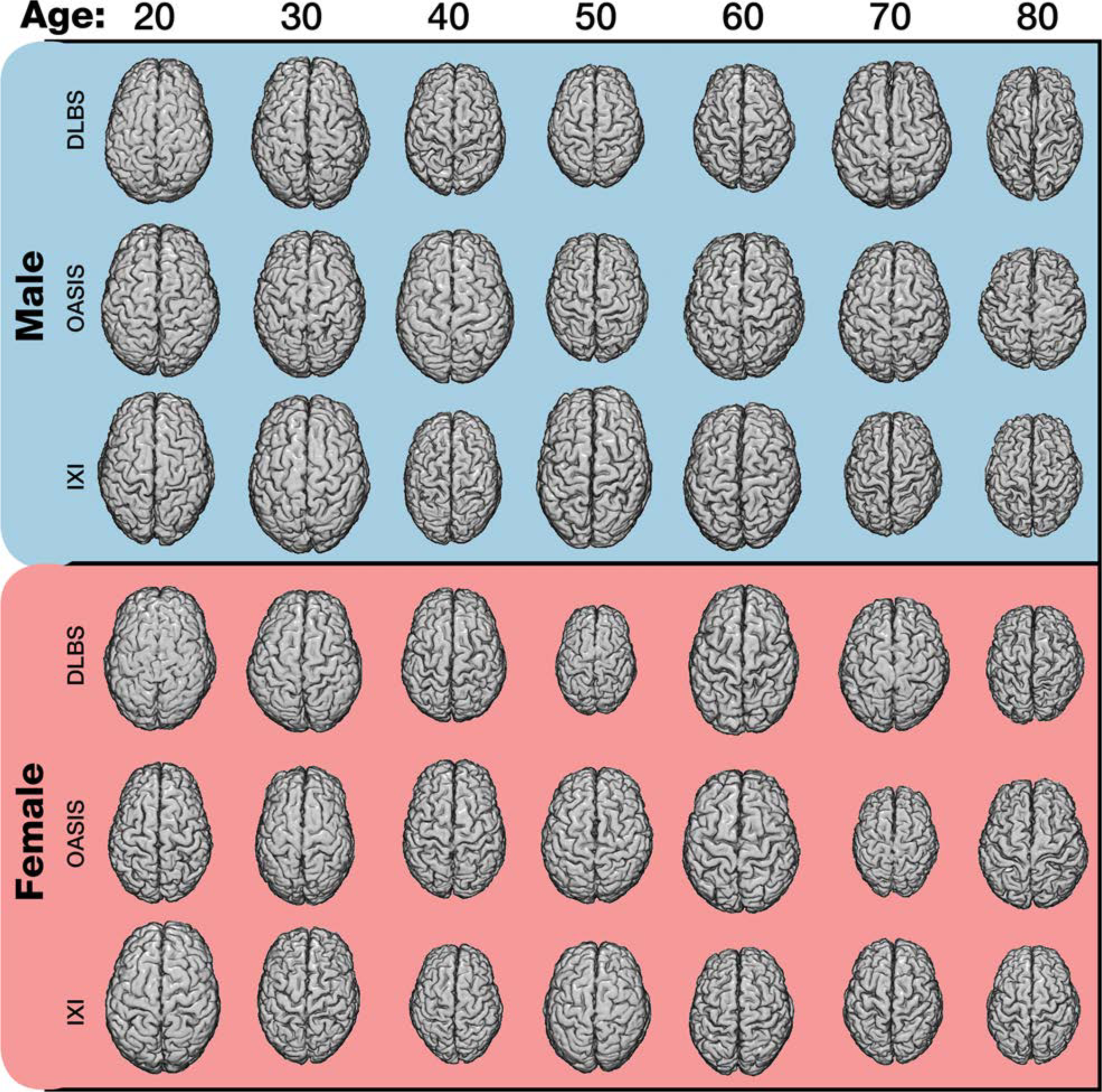
Cortical surfaces for each age decade, sex, and training sample. Representative cortical surface reconstructions for individuals with ages in the first year of each decade (with the exception of 40s, where there were insufficient male participants between ages 40 and 41). All reconstructions are shown at the same scale. 3D reconstruction images were generated as described in Madan (2015).

Here we sought to predict an individual’s age from their brain morphology, attempting to optimize the brain parcellation and segmentation techniques so as to maximize their predictive accuracy. Though a few others have similarly sought to predict an individual’s age from structural MRI volumes (e.g., Ashburner, 2007; Cole et al., 2015; Franke & Gaser, 2012; Franke et al., 2010, 2014; Schnack et al., 2016; see Cole and Franke, 2017, for a review), these implementations relied on voxel-based morphometry (VBM) rather than regional surface-based morphology estimates and thus cannot be used to assess the predictive value of *specific* morphological features (e.g., thickness, gyrification) and cortical granularity as sought here. To construct this age-prediction model, we used several open-access MRI datasets. The public sharing of MRI data has been quickly growing and a large number of open-access datasets are now available (Biswal et al., 2010; Das et al., 2017; Madan, 2017; Mennes et al., 2013; Poldrack & Gorgolewski, 2014; Poldrack et al., 2017; Shenkin et al., 2017). The use of open-access datasets enabled us to have the large sample sizes needed to have training datasets as well as held-out datasets, as well as demonstrate the generalizability and reproducibility of the presented results, though this was not possible only a few years ago (Dickie et al., 2012).

Figure 3 provides an overview of factors that are known to influence estimates of brain morphology. These factors can be quite varied, ranging from transient changes, such as time-of-day (Nakamura et al., 2015; Trefler et al., 2016) and hydration (Duning et al., 2005; Nakamura et al., 2014), to more long-lasting changes, such as exercise (Hayes et al., 2014; Steffener et al., 2016), diet/lifestyle (Booth et al., 2015; Khan et al., 2015; Kullmann et al., 2016), and meditation (Tang et al., 2015). Brain morphology has also been linked with genetic variations, such as *APOE* and *BDNF* (see Strike et al., 2015, for a comprehensive review). As such, it is important to acknowledge the breadth of effects that influence any measure of brain morphology. Although some of these sources of variability would be minimized if multiple scans were taken (e.g., at different times of day or with different levels of hydration), any model predicting an individual’s age from *only* brain morphology estimates will be unable to account for some additional sources of variance and thus will have some degree of error. Nevertheless, it is useful to understand how well age predictions can be made on the basis of a structural scan that can be acquired in just a few minutes. Not only is this a relevant exercise for confirming the aspects of brain structure that are most strongly associated with age-associated differences, it also has potential clinical relevance; if the brain structure of healthy adults is a reasonable predictor of their age, then failures in age prediction (e.g., a structural scan that suggests someone is a decade older than they are) may help to indicate the presence of a prodromal state (Cole et al., 2015; Franke & Gaser, 2012; Franke et al., 2010, 2014; Schnack et al., 2016; see Cole & Franke, 2017, for a review). While the current focus is on age-related differences through the adult lifespan, brain structure has also been examined through development (Brown et al., 2012; Dosenbach et al., 2010; Lee et al., 2014; Mills et al., 2016; Qin et al., 2015; Somerville, 2016).

**Figure 3.**
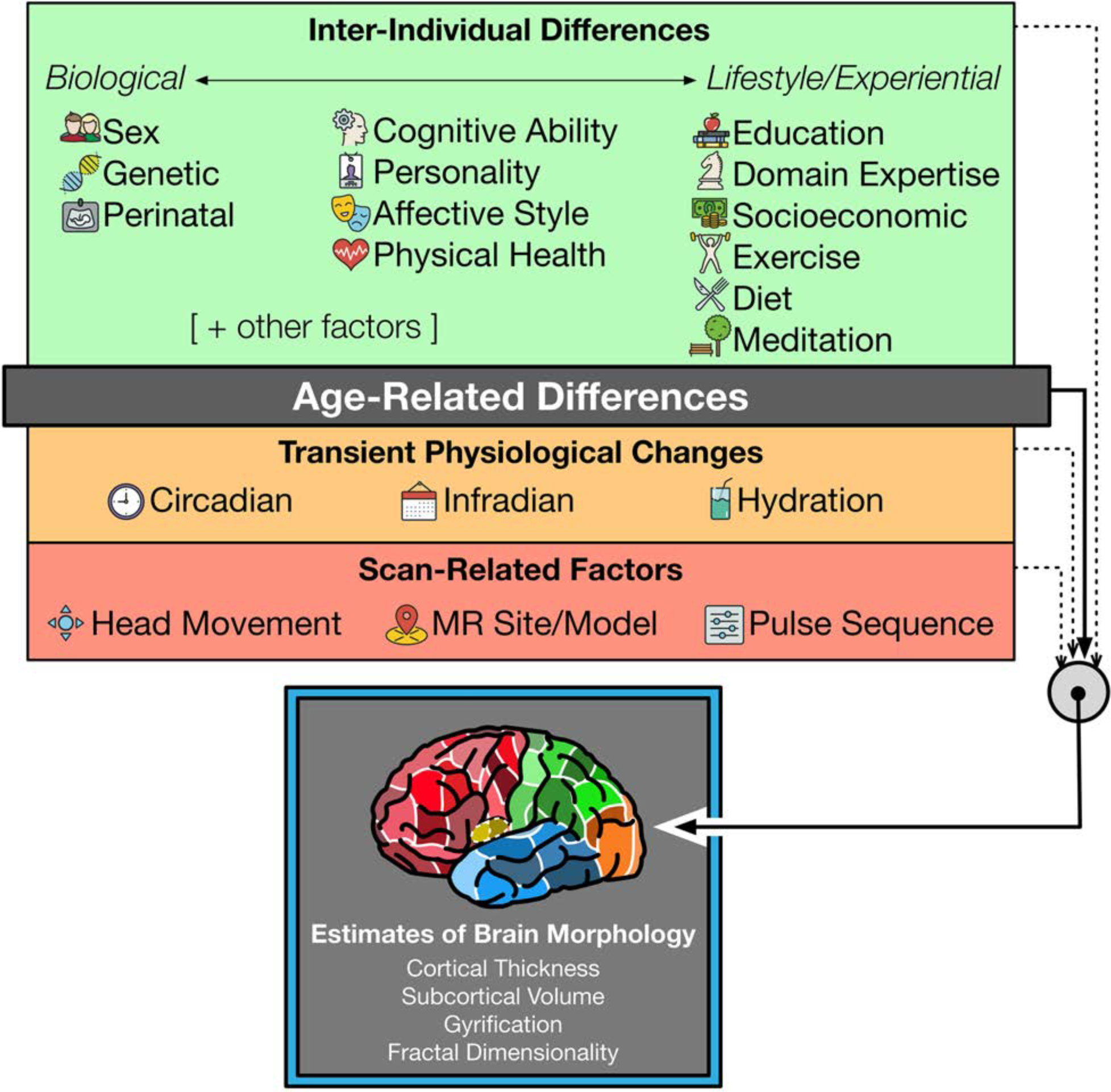
Overview of factors known to influence estimates of brain morphology.

## Procedure

### Datasets

Three datasets were used to train the age-prediction algorithm, with an additional two datasets used as independent test samples. The age distribution for each of the datasets are shown in Figure 4.

**Figure 4.**
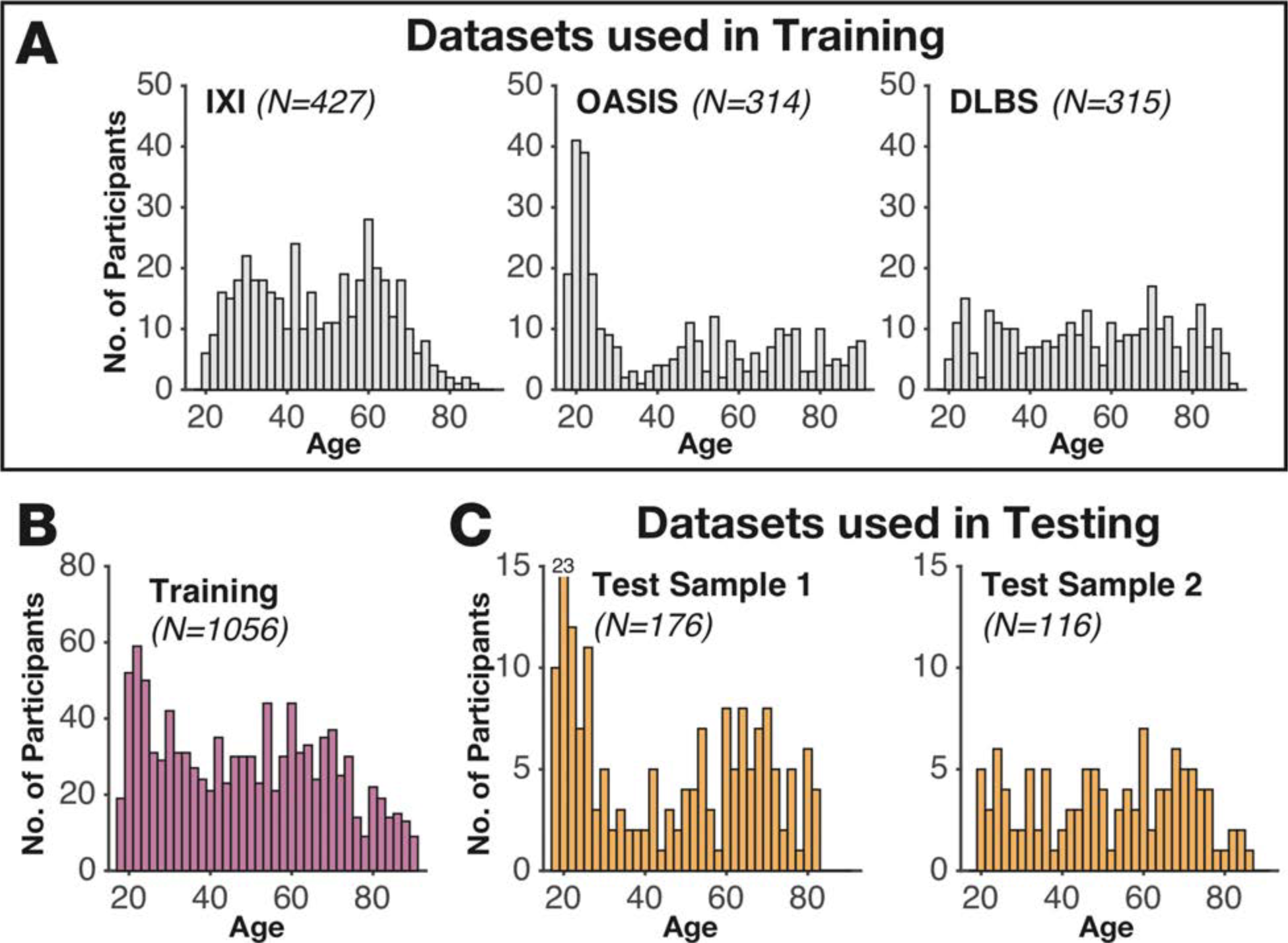
Age distributions for each of the datasets. (A) Age distributions for each of the datasets used in the training. (B) Age distribution for the aggregated training dataset (i.e., combining IXI, OASIS, and DLBS). (C) Age distributions for the independent test datasets.

#### Training Sample 1 (IXI)

consisted of 427 healthy adults, (260 females) aged 20-86, from the publicly available Information extraction from Images (IXI) dataset (http://brain-development.org/ixi-dataset/). This is the same subset of individuals we used previously to investigate age-related differences in cortical regions; see Madan and Kensinger (2016, 2017a) for further details.

#### Training Sample 2 (OASIS)

consisted of 314 healthy adults (196 females), aged 18-94, from the publicly available Open Access Series of Imaging Studies (OASIS) cross-sectional dataset (Marcus et al., 2007; http://www.oasis-brains.org). Participants were screened for neurological and psychiatric issues; the Mini-Mental State Examination (MMSE) and Clinical Dementia Rating (CDR) were administered to participants aged 60 and older. In the current sample, participants with a CDR above zero were excluded; all remaining participants scored 25 or above on the MMSE. Multiple T1 volumes were acquired using a Siemens Vision 1.5 T with a MPRAGE sequence; only the first volume was used here. Scan parameters were: TR=9.7 ms; TE=4.0 ms; flip angle=10°; voxel size=1.25×1×1 mm. This sample was previously used in Madan and Kensinger (2017a) and Madan (in press).

#### Training Sample 3 (DLBS)

consisted of 315 healthy adults (198 females), aged 20-89, from wave 1 of the Dallas Lifespan Brain Study (DLBS), made available through the International Neuroimaging Data-sharing Initiative (INDI; Mennes et al., 2013) and hosted on the the Neuroimaging Informatics Tools and Resources Clearinghouse (NITRC; Kennedy et al., 2016) (http://fcon1000.projects.nitrc.org/indi/retro/dlbs.html). Participants were screened for neurological and psychiatric issues. All participants scored 26 or above on the MMSE. T1 volumes were acquired using a Philips Achieva 3 T with a MPRAGE sequence. Scan parameters were: TR=8.1 ms; TE=3.7 ms; flip angle=12°; voxel size=1×1×1 mm. See Kennedy et al. (2015) and Chan et al. (2014) for further details about the dataset; this sample was previously used in Madan (in press).

#### Test Sample 1

consisted of 176 healthy adults (89 females), aged 18-83, recruited by the Cognitive and Affective Laboratory at Boston College (BC) and screened for neurological and psychiatric issues, and to have scored above 26 on the MMSE. T1 volumes were acquired using a Siemens Trio 3 T with a MEMPRAGE sequence optimized for morphometry studies (van der Kouwe et al., 2008; Wonderlick, et al., 2009). Scan parameters were: TR=2530 ms; TE=1.64, 3.50, 5.36, 7.22 ms; flip angle=7°; voxel size=1×1×1 mm. This sample was previously used in Madan and Kensinger (2017a).

#### Test Sample 2

consisted of 116 healthy adults (70 females), aged 20-87, recruited by Dr. Craig Stark’s laboratory at University of California–Irvine (UCI) and screened for neurological and psychiatric issues, and to have scored above 26 on the MMSE, made available on NITRC (Kennedy et al., 2016) (https://www.nitrc.org/proiects/stark_aging/). T1 volumes were acquired using a Philips Achieva 3 T with a MPRAGE sequence. Scan parameters were: TR=11 ms; TE=4.6 ms; flip angle=12°; voxel size=0.75×0.75×0.75 mm. See Stark et al. (2013) and Bennett et al. (2015) for further details about the dataset.

### Pre-processing of the structural MRIs

Data were analysed using FreeSurfer v.5.3.0 (https://surfer.nmr.mgh.harvard.edu) on a machine running CentOS 6.6. FreeSurfer was used to automatically volumetrically segment and parcellate cortex from the T1-weighted images (Dale et al., 1999; Fischl, 2012; Fischl & Dale, 2000; Fischl et al., 1999, 2002, 2004). FreeSurfer’s standard pipeline was used (i.e., `recon-all`) and no manual edits were made to the surface meshes. Cortical thickness is calculated as the distance between the white matter surface (white-gray interface) and pial surface (gray-CSF interface) (Dale et al., 1999; Fischl & Dale, 2000). Thickness estimates have previously been found to be in agreement with manual measurements from MRI images (Kuperberg et al., 2003; Salat et al., 2004), as well as *ex vivo* tissue measurements (Cardinale et al., 2014; Rosas et al., 2002). Gyrification was calculated using FreeSurfer, as described in Schaer et al. (2012).

#### Cortical parcellations

Here we used seven parcellation atlases to determine the amount of relevant age-related differences in cortical structure: (1) entire cortical ribbon (1 region; i.e., unparcellated); (2) each of the four lobes (4 regions); (3) Desikan-Killany-Tourville (DKT) atlas (62 regions; Klein & Tourville, 2012); (4) Destrieux et al. (2010) atlas (148 regions); (5) von Economo-Koskinas atlas (86 regions; Scholtens et al., 2018); (6) Brainnetome atlas (210 regions; Fan et al., 2016); and (7) Lausanne atlas (1000 regions; Hagmann et al., 2008). Each of these parcellation approaches is shown in Figure 1. The DKT and Destrieux atlases are included as standard parcellation atlases within FreeSurfer. The lobe parcellation was delineated by grouping parcellation regions from the Destrieux atlas, as done in Madan and Kensinger (2016).

The von Economo-Koskinas, Brainnetome, and Lausanne atlases were applied to each individual’s reconstructed cortical surface using `mris_ca_label` with the cortical parcellation atlas files (`*.gcs`) that have been distributed online by the respective researchers. The von Economo-Koskinas atlas was implemented in FreeSurfer by Scholtens et al. (2018; http://www.dutchconnectomelab.nl/economo/), based on the cortical areas identified by von Economo and Koskinas (1925). The Brainnetome atlas was developed by Fan et al. (2016; http://atlas.brainnetome.org) based on a modified Desikan-Killany atlas (Desikan et al., 2006; a precursor to the DKT atlas) that initially reduced the number of cortical regions to 20 per hemisphere (rather than the 34), but then further parcellated the cortex using connectivity data extracted from diffusion and resting-state scans, resulting in a total of 210 regions. The Lausanne atlas was similarly initially constructed using the Desikan-Killany atlas, then further subdivided each region into smaller patches of approximately equivalent area (Hagmann et al., 2008). The Lausanne atlas is distributed as part of the Connectome Mapper (Daducci et al., 2012; https://github.com/LTS5/cmp).

#### Fractal dimensionality

The complexity of each structure was calculated using as the fractal dimensionality of the filled structure. Our work previously demonstrated that fractal dimensionality indexes age-related differences in cortical and subcortical structures better than extant measures (i.e., cortical thickness, cortical gyrification, subcortical volume), where older adults exhibit reductions in structural complexity relative to younger adults (Madan & Kensinger, 2016, 2017a; Madan, in press). In fractal geometry, several approaches have been proposed to quantify the ‘dimensionality’ or complexity of a fractal; the approach here calculates the Minkowski–Bouligand or Hausdorff dimension (see Mandelbrot, 1967). This structural property can be measured by considering the 3D structure within a grid space and counting the number of boxes that overlap with the edge of the structure. By then using another grid size (i.e., changing the box width), the relationship between the grid size and number of counted boxes can be determined (‘box-counting algorithm’). Here we used box sizes (in mm) corresponding to powers of 2, ranging from 0 to 4 (i.e., 2^*k*^ [where *k* = {0, 1, 2, 3, 4}] = 1, 2, 4, 8, 16 mm). The slope of this relationship in log-log space is the fractal dimensionality of the structure. Thus, the corresponding equation is:

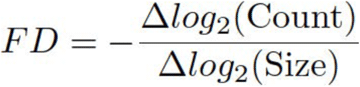

If only the boxes overlapping with the edge/surface of the structure are counted, this slope represents the fractal dimensionality of the *surface* (*FD*_s_). If the boxes within the structure are additionally counted, the resulting slope represents the fractal dimensionality of the *filled* volume (*FD*_*f*_; see Figure 5). As the relative alignment of the grid space and the structure can influence the obtained fractal dimensionality value using the box-counting algorithm, we instead used a dilation algorithm that is equivalent to using a sliding grid space and calculating the fractal dimensionality at each alignment (Madan & Kensinger, 2016, 2017b).

**Figure 5.**
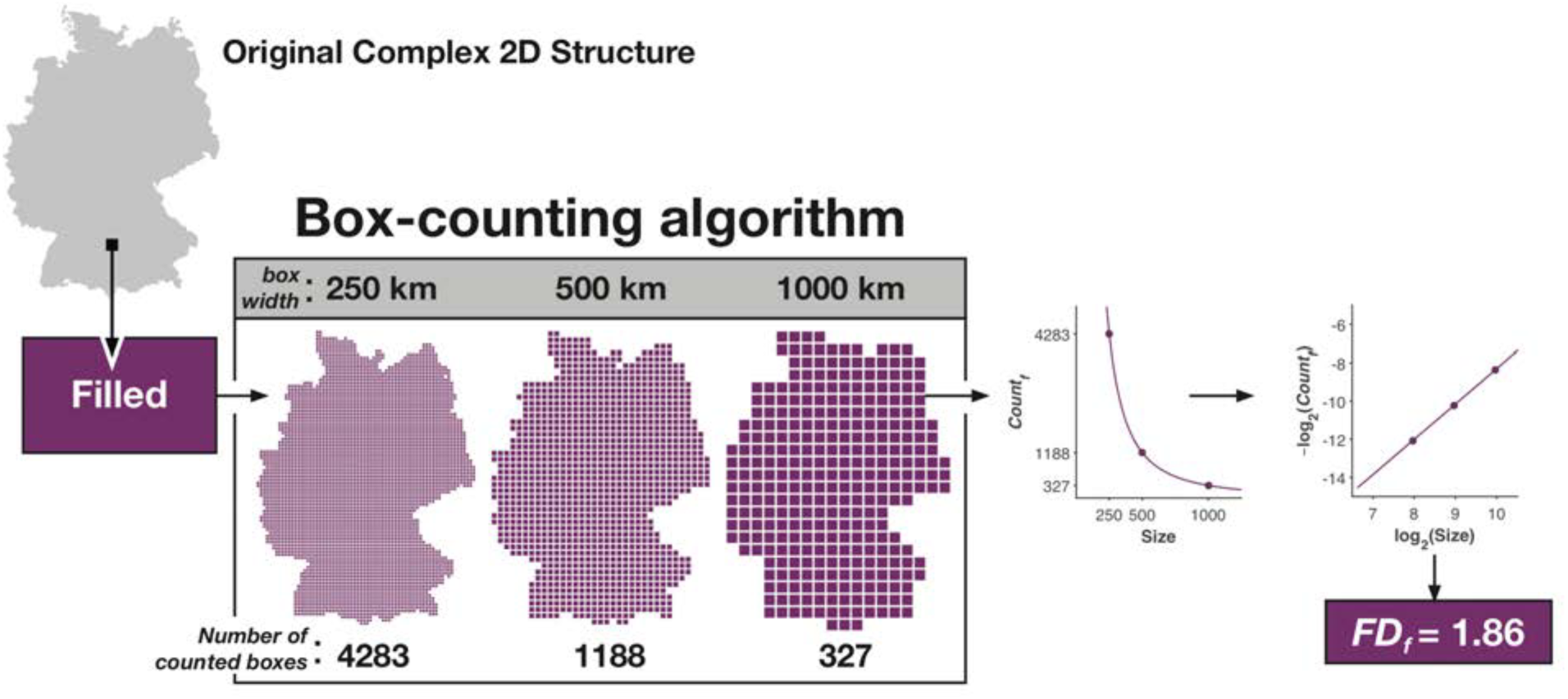
Illustration of how fractal dimensionality is measured from a 2D structure. Modified from Madan and Kensinger (2016).

Fractal dimensionality was calculated using the calcFD toolbox (Madan & Kensinger, 2016; http://cmadan.github.io/calcFD/). Briefly, the toolbox calculates the ‘fractal dimensionality’ of a 3D structure, and is designed to use intermediate files from the standard FreeSurfer analysis pipeline. calcFD was first applied to measure the complexity of the cortical ribbon and lobes (Madan & Kensinger, 2016), but has been extended to subcortical structures (Madan & Kensinger, 2017a; Madan, in press), and the DKT parcellation atlas (Madan & Kensinger, 2017b).

Here we only used the fractal dimensionality of the filled structures (*FD*_*f*_), as this measure has been demonstrated to be more sensitive, than the fractal dimensionality of the surface, to age-related differences in cortical structure (Madan & Kensinger, 2016). For each participant, the fractal dimensionality was calculated for: (a) entire cortical ribbon (1 region); (b) four lobes (4 regions); (c) DKT-parcellated atlas (62 regions); and (d) von Economo-Koskinas atlas (86 regions). As smaller parcellations of cortex inherently have decreased fractal dimensionality, i.e., becoming closer to a truncated rectangular pyramid, we did not calculate the fractal dimensionality for the parcellation atlases with smaller regions (see Figure 1) (Madan & Kensinger, 2017a, 2017b). Indeed, prior analyses indicated that the parcellations in the Destrieux atlas were too fine-grained for meaningful fractal dimensionality calculations, but that the regions within the DKT atlas were of sufficient size (Madan & Kensinger, 2017b).

### Age-prediction algorithm

A machine learning approach was used to predict age (in years) from brain morphology. Specifically, we trained a machine-learning regression framework with the aggregated training data (*N*=1056), using brain morphology data to predict the age data. We then applied the trained regression model to the brain morphology data from the test samples to obtain age predictions. Age predictions were then compared with the actual age data. Performance of the age-predictions was evaluated using two metrics: *R*^*2*^ and *MdAE*. *R*^*2*^ was used as a measure of explained variability. Median absolute error (*MdAE*) has been found to be more robust to outliers than mean squared error (*MSE*) (Armstrong & Collopy, 1992; Hyndman & Koehler, 2006).

The machine-learning regression framework primarily relies on several statistical methods to minimize over-fitting through regularization, dimensionality reduction, and feature selection. This is accomplished through sequentially applying three statistical techniques: smoothing spline regression, PCA, and relevance vector regression (RVR)—which we have implemented as a MATLAB toolbox called Prism (Madan, 2016). These statistical techniques are described further below. As a whole, Prism is a MATLAB toolbox designed for conducting spline-based multiple regression using training and test datasets. Benchmark analyses indicated that Prism provided better age-prediction estimates than standard RVR (Madan, 2016). Prism was configured to (a) use least-squares splines were to reduce over-fitting (smoothness parameter set to 0.1) and (b) keep the principal components that explained the most variance, retaining as many components as necessary to explain 95% of the variance. Values were *Z*-scored prior to the smoothing spline regression to ensure that different measures were smoothed to the same degree, as the smoothness parameter is influenced by the overall magnitude of the input values (as in Madan, in press).

#### Relevance Vector Regression (RVR)

To make predictions using the structural measures, we employed relevance vector regression (RVR). RVR is the application of relevance vector machine or RVM to a regression problem, and is a relatively recent machine learning technique (Tipping, 2000, 2001; Tipping & Faul, 2003). In functionality, RVM shares many characteristics with support vector machines (SVM), but is generally more flexible (Tipping, 2000). RVM can also be considered as a special case of a Sparse Bayesian framework (Tipping, 2001; Tipping & Faul, 2003) or Gaussian process (Rasmussen & Williams, 2006). For a more in-depth discussion of RVM, see Bishop (2006).

Broadly, RVR is similar to multiple linear regression with regularization (e.g., LASSO and ridge regression; see Hastie et al., 2009; Tibshirani, 1996), which reduces the number of predictors/model complexity by removing those predictors that are redundant, reducing over-fitting and yielding a more generalizable model. The implementation of this procedure is sometimes referred to as automatic relevance determination (ARD) (MacKay, 1996; Wipf & Nagarajan, 2007). Further, it has been suggested that RVR is comparable to a Bayesian implementation of LASSO regression (Gao et al., 2010; Jamil & ter Braak, 2012; Park &

Casella, 2008). Here we used the MATLAB implementation of RVM that uses an accelerated training algorithm (Tipping & Faul, 2003), freely available from the author’s website (http://www.miketipping.com/sparsebayes.htm).

#### Smoothing spline regression

While age-related changes in morphology are often modeled using linear and quadratic functions (e.g., Hogstrom et al., 2013; Madan & Kensinger, 2016, Sowell et al., 2003; Walhovd et al., 2011), non-linear functions have been shown to better model age-related differences (Fjell et al., 2010, 2013). This approach uses smoothing spline regression, where the relationship between a predictor and dependent variable is fit using piece-wise cubic functions (Fox, 2000; Wahba & Wold, 1975; Wahba, 1990). Cubic smoothing spline is implemented in MATLAB as the function `csaps`. Here each predictor (i.e., brain morphology measure) was treated as an independent predictor of age and were individually regressed versus age. To combine these estimates, i.e., for multiple regression, the Prism toolbox was used (Madan, 2016).

#### Held-out test data

Age prediction performance here was evaluated using independent test datasets. This approach was taken to ensure that the age predictions were not biased—for instance, numerous studies have demonstrated that performance with *k*-fold cross-validation can lead to over-fitting in the model selection (e.g., Golbraikh & Tropsha, 2002; Rao et al., 2008; Saeb et al., 2016; Skocik et al., 2016).

#### Pre-and post-processing

Data were maintained as three separate datasets: Training, Test Sample 1, and Test Sample 2. Within each dataset, main effects of sex were initially regressed out before being entered into the machine-learning regression framework; the main effect of site was also regressed out for the Training dataset. Outputted age predictions were mean centered (to the mean of the predicted ages) to compensate for sample differences (as in Franke et al., 2010; Franke & Gaser, 2012). The variance in the predicted ages in the test data was also matched to variance of the training data, to correct for a regression-to-the-mean bias.

## Results

### Regional age-related differences in cortical structure

In our prior work investigating age-related differences in cortical structure (Madan & Kensinger, 2016), we had only used bilateral lobe parcellations (i.e., 4 regions). However, in subsequent work, our fractal dimensionality analyses were used with the DKT parcellation, with 62 cortical parcellations (Madan & Kensinger, 2017b). While we use this improved parcellation method in the current paper, along with multiple regression methods to estimate age predictions in held-out test samples, we thought it would be useful to calculate the relationship between structural measures for each parcellation with age.

We used a smoothing spline regression approach, as described in the methods, with the full training dataset (*N*=1056). All of the regressions controlled for effects of site.

We first conducted smoothing-spline regressions using the cortical ribbon measures. Inter-individual differences in mean cortical thickness explained 47.17% of the variance in age (i.e., *R*^*2*^). Mean gyrification explained 36.07% of the variance; fractal dimensionality (of the filled volume) explained 73.53% of the variance. Reassuringly, these values correspond reasonably well with the variances explained in the independent, held-out test samples (see Table 1). For comparison, with linear and quadratic effects in the IXI sample alone, these values were 33.55%, 20.61%, and 51.66%, respectively (as reported in Madan & Kensinger, 2016); the relative increases in the explained variance correspond to the use of spline regression, instead of linear and quadratic regression, and also indicate that our overall approach and between-site harmonization was successful.

**Table 1.**
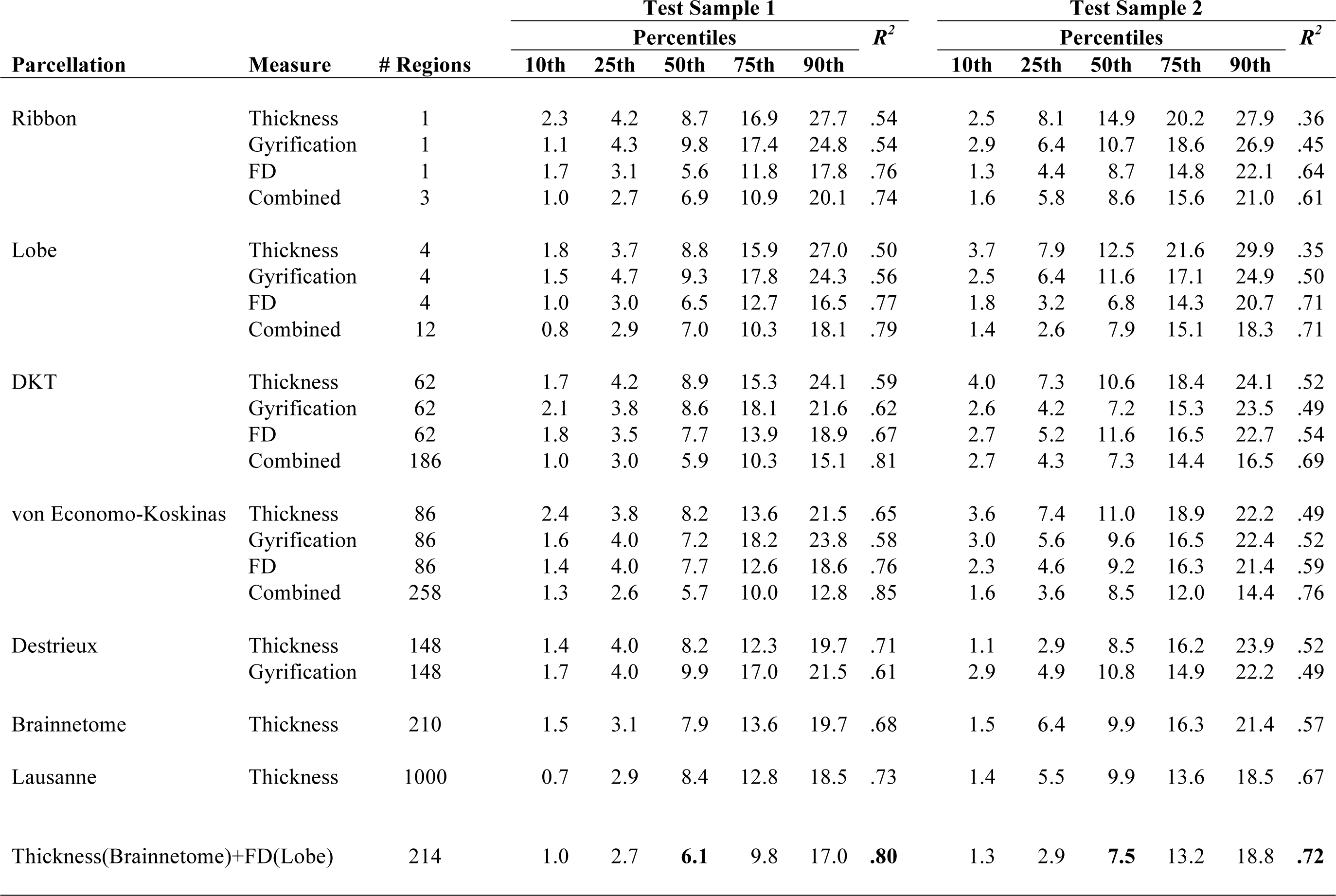
Summary of age-prediction model performance.

We subsequently conducted similar smoothing-spline regressions for each of the 62 cortical parcellations of the DKT atlas, as shown in Figure 6A. The cortical thickness and gyrification analyses correspond well with the comparable analysis by Hogstrom et al. (2013). Age-related differences in cortical thickness are most pronounced in frontal regions and the superior temporal gyrus, and are least effected in occipital regions, the superior parietal lobule, and the postcentral gyrus. In contrast, age-related differences in gyrification are most pronounced in the pre‐ and postcentral gyri, caudal middle frontal gyrus, as well as supramaginal gyrus and inferior parietal lobule. Frontal regions and parcellations on the medial surface have the least age-related differences in gyrification.

**Figure 6.**
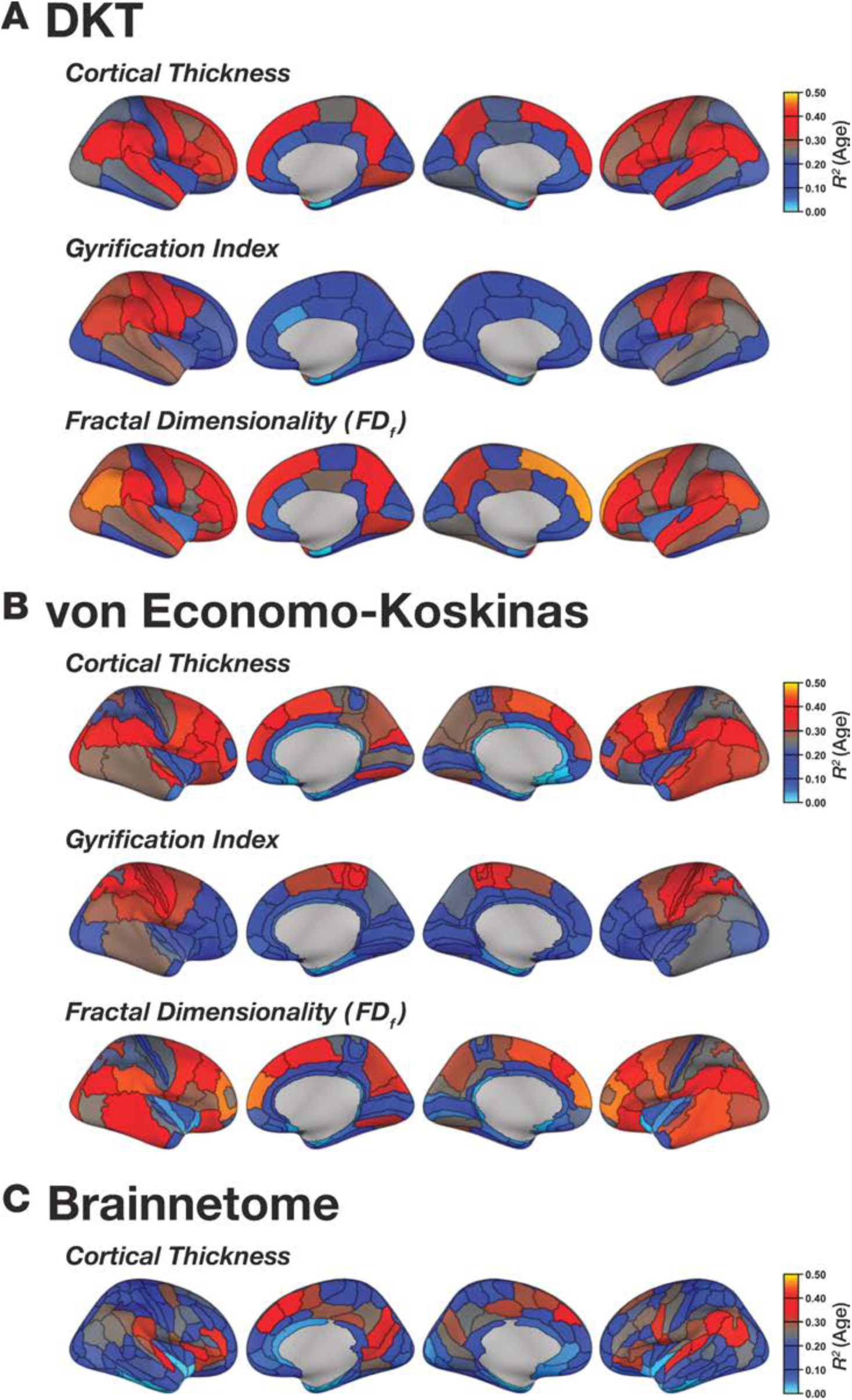
Regional age-prediction differences (*R*^*2*^) using the (A) DKT, (B) von Economo-Koskinas atlases, and (C) Brainnetome for the respective structural measures using smoothing-spline regression.

Age-related differences in fractal dimensionality have a topology that resembles a mixture of the thickness and gyrification patterns, but is generally higher in explained variance (i.e., *R*^*2*^). Regional fractal dimensionality was most affected by age in the superior temporal gyrus and inferior parietal lobule, though frontal regions were also markedly affected. Inter-individual differences in fractal dimensionality of the inferior temporal gyrus and regions along the medial surface were the least associated with age.

As shown in Figure 6B, we also conducted a similar analysis using the von Economo-Koskinas atlas (Scholtens et al., 2018) and found a similar pattern of results across the parcellations and cortical measures. Figure 6C shows that the most age-sensitive regions from the other atlases generalize to the Brainnetome atlas as well, however, thickness of many parcellations are only weakly related to age, likely due to the substantially smaller regions here and an increasing role of inter-individual differences and regional specificity.

### Age-prediction models

To investigate the relationship between brain morphology and age-prediction performance, we first calculated a benchmark measure, then compared age-prediction models using regional cortical thickness for each of the seven parcellation approaches. Next we compared different structural measures (i.e., thickness, gyrification, fractal dimensionality, and a combination of all three) across four parcellation atlases. Finally, we took a theoretical approach to combining the morphological measures to best capture age-related differences in structure. Results from the age-prediction models are summarized in Table 1.

#### Benchmark performance

To provide a benchmark to evaluate the age-prediction performance in the held-out test data using the various estimates of brain morphology, a simple estimation of a lower-bound (shown in light gray in Figure 7) was determined by ‘predicting’ that all participants’ ages in the test dataset were the mean of the dataset. As such, *R*^*2*^=0 for this benchmark model, and we would not expect any reasonable age-prediction model to produce worse estimates than this.

**Figure 7.**
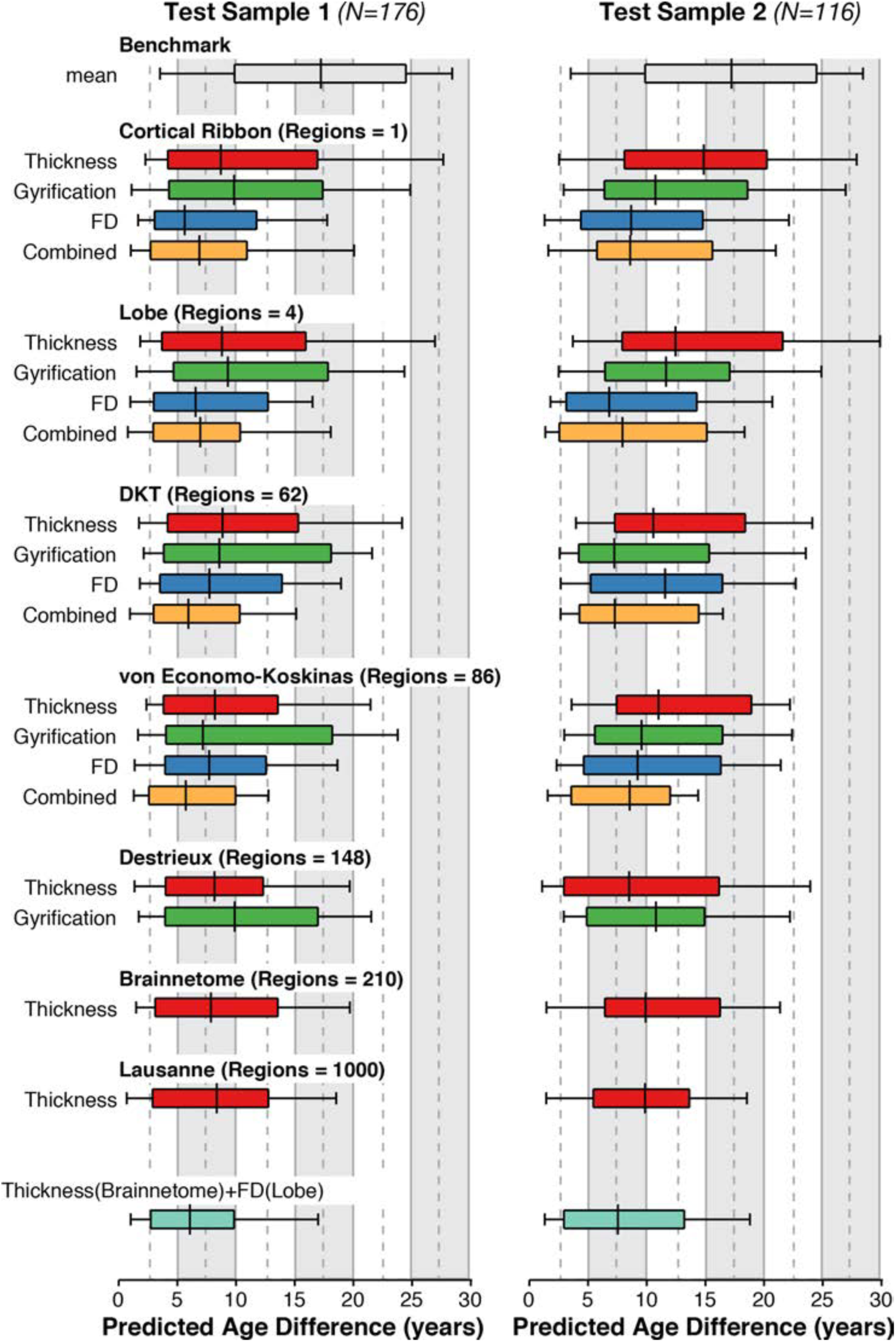
Age-prediction performance for the models based on each of parcellation atlases and structural measures. ‘*Regions*’ corresponds to the number of regions/predictors used in the model (see Figure 1). ‘*FD*’ denotes fractal dimensionality of the filled structures. Each box-and-whisker bar denotes the median (i.e., *MdAE*) as a tick-mark. The box spans the 25th to 75th percentiles; the whiskers span the 10th to 90th percentiles. Values for each of these percentiles, as well as *R*^*2*^, are reported in Table 1.

In Figure 7, each box-and-whisker denotes the median (i.e., *MdAE*) as a tick-mark. The box spans the 25th to 75th percentiles; the whiskers span the 10th to 90th percentiles.

#### Cortical thickness

Performance for the age-prediction models based on cortical thickness, for each of the seven parcellation approaches, are shown in Figure 7 and Table 1. Results show that fmer-grain parcellations generally yielded better age predictions, demonstrating that regional heterogeneity in age-related differences in cortical thickness led to improved prediction performance. Age-prediction models using the cortical ribbon and lobe parcellations performed comparably, indicating that the slight increase in heterogeneity with the lobe parcellations was not sufficient to improve predictions. However, performance on the atlases that had more fine-grain parcellations, led to a decrease in the median absolute error of approximately three years. Performance between these different atlases did not markedly differ, with age predictions having a median error of 7 to 10 years. While performance with the Lausanne atlas (which comprised 1000 regions) was relatively good, it was not better than the parcellation atlases that had far fewer regions. Here we considered the Brainnetome atlas to be the best performing model based on cortical thickness estimates.

#### Comparing different measures of morphology

Next, we compared age-prediction models based in cortical thickness, gyrification, and fractal dimensionality. As indicated earlier, fractal dimensionality inherently becomes less sensitive with smaller parcellations and was only used for the cortical parcellations with less than 100 regions.

Beginning with models using only a single value for the unparcellated cortical ribbon, the model based on the fractal dimensionality did markedly better than those based on mean cortical thickness or gyrification. Combining the three measures (i.e., three predictors in the regression) led to worse age predictions than complexity alone, due to insufficient feature selection and/or possible over-fitting. Given that there were only three predictors input, this suggests that the automatic relevance determination used in RVR was not ‘strict’ enough to only use complexity, though later combined models were able to perform better than their component models that relied on only a single measure of morphology. Performance on the age-prediction models using the lobe-wise parcellation scheme performed relatively similarly to those that only used the cortical ribbon.

The models that used the DKT parcellation atlas using only one of the morphology measures performed comparably to those with the unparcellated ribbon and lobe parcellations, however, the combined model here did perform notably better. The combined model had a median error of 8 years and *R*^*2*^ values near .70. As noted earlier, age predictions based on cortical thickness improved markedly when using the DKT parcellation, relative to the ribbon and lobe parcellations. Importantly, we also found that the fractal dimensionality model using the DKT parcellation was performing worse than the ribbon and lobe models, suggesting that using the finer-grain parcellations was resulting to notable losses in complexity-related information, as alluded to previously. Interestingly, the age-prediction models based on the von Economo-Koskinas atlas performed comparably with the DKT atlas when each structural metric was used independently, but error was sufficiently lower when the metrics were combined.

#### Combining measures

The four combined models are shown in orange in Figure 7. Based on the performance of these age-prediction models, we constructed an additional model, combining the two best cortical measures, across different parcellation approaches—cortical thickness from the regions of the Brainnetome parcellation atlas, and fractal dimensionality from the lobe-wise parcellation. This newly constructed model (shown in light green) out-performed the component models, indicating that the two structural measures each provided unique age-related difference characteristics. A scatter plot of the predicted and actual ages for this model is shown in Figure 8.

**Figure 8.**
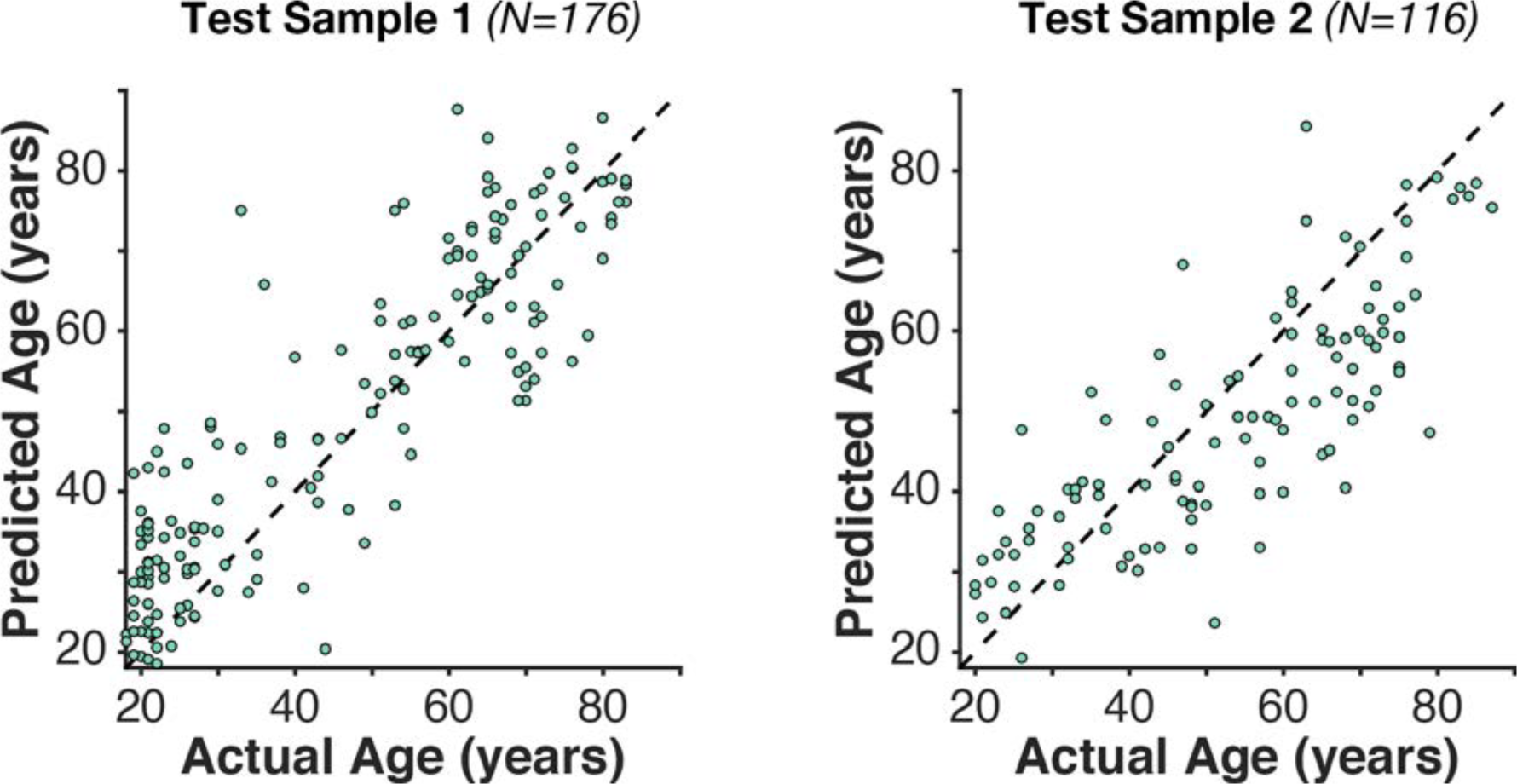
Scatter plot of the actual and predicted age for the best-fitting model.

## Discussion

The present study reveals that inter-individual differences in cortical structure are not only strongly correlated with age, but also can robustly be used to predict age. Although almost any metric of cortical structure can predict age to within a decade (10-12 years), the present study also reveals that by optimizing metrics, a much more accurate prediction (6-7 years) can be achieved. In-line with prior results (Hogstrom et al., 2013; Madan & Kensinger, 2016), we found that differences in gyrification are not as related to age as cortical thickness, and fractal dimensionality was a better morphological metric of age-related differences than either of these other metrics. Importantly, the best age predictions were achieved when both fractal dimensionality and cortical thickness were combined, demonstrating that these metrics contribute unique information about age-related differences in brain structure.

Across different cortical parcellation schemes, cortical thickness estimates localized to different gyri and sulci are substantially more indicative of age-related differences relative to lobe-wise measurement. The recently developed Brainnetome atlas was selected as the best atlas for representing age-related differences in cortical thickness, though the DKT, Destrieux, and von Economo-Koskinas atlases did not markedly differ in performance. Parcellating the cortex into even more constrained patches (i.e., the Lausanne atlas) did not provide any additional predictive value, indicating that the level morphological granularity in the Brainnetome parcellation was sufficient for age predictions based on cortical thickness estimates. Fractal dimensionality inherently becomes less sensitive with smaller parcellations, and performed best when only using lobe-wise parcellations, rather than the more fine-grained parcellations of the DKT and von Economo-Koskinas atlases. Gyrification did not appear to be sufficiently predictive of age to further contribute to predictions. Without pooling across different atlases, the von Economo-Koskinas atlas produced relatively good age predictions when the three cortical metrics were combined.

Within the broader literature, the age-prediction approach here can serve as a baseline model of healthy, successful aging. For instance, this approach can be used to characterize which regions exhibit more or less age-related variability. Since the training and test datasets were wholly independent, age-predictions applied to additional independent datasets should be indicative of healthy aging. Future studies could use the current approach with smaller samples, where larger differences between predicted age and chronological age would be indicative of cohort characteristics related to atypical aging. Of course, it would still be optimal to have MRI data for ‘healthy’ individuals acquired alongside the group of interest to account for differences related to scanner hardware, pulse sequence, or potential differences in demographics (e.g., years of education).

Although the present results reveal systematic age-related differences in brain structure, several questions remain about the nature of these age differences. For instance, the cortical measures used here are limited to macroscopic differences, as differences in the microstructure (e.g., neuron density, neuropil composition) are beyond the capabilities of current MRI methods. This raises the question of what neurobiological changes are manifesting in these macroscopic changes in structure. In a different vein, in models where age-predictions performed relatively poorer, and in light of the myriad of ways that individual brains can differ, one must wonder what other non-aging factors may be manifesting in morphological differences and minimizing the ability to isolate age differences. In the following two sections we outline the extant literature that investigates these two topics.

### Neurobiological basis of age-related differences in brain structure

While a large number of studies examining age-related differences in cortical thickness have found reduced cortical thickness in older adults, particularly in frontal regions, the underlying changes in the cortical microstructure are unclear (Dickstien et al., 2013; Koo et al., 2012; Meyer et al., 2014; Sowell et al., 2004). One reason this has yet to be resolved is that to do so appropriately would require not only MRI data, but also post-mortem histological tissue samples. Extant histological evidence suggests that age related differences in cortical regions is generally not associated with a decrease in the number or size of neurons (Freeman et al., 2008; Gefen et al., 2015; Morrison & Hof, 1997; Peters & Rosene, 2003; Peters et al., 1998; Uylings & de Brabander, 2002; von Bartheld, in press), despite initial suggestions to the contrary (Terry et al., 1987). What does appear to differ, however, is the dendritic structure, particularly in pyramidal neurons, as well as other features of the neuropil (Casanova et al., 2011; Dickstein et al., 2007, 2013; Duan et al., 2003; Eickoff et al., 2005; Hao et al., 2007; Morrison & Baxter, 2012; Peters, 2002; Scheibel et al., 1975; Uylings & de Brabander, 2002). Convergently, pyramidal neurons are particularly prominent in prefrontal cortex, as well as prefrontal pyramidal neurons having orders of magnitude more dendritic spines than those in some other regions (Elston, 2003).

The literature investigating the biological basis for age-related differences in gyrification is even more sparse. Unlike cortical thickness, age-related differences in gyrification are most pronounced in the parietal lobe (Hogstrom et al., 2013; Madan & Kensinger, 2016). In a large sample of chimpanzees, Autrey et al. (2014) observed increased gyrification in adults relative to adolescents, consistent with human data; however, an age-related decrease in gyrification was not observed. The authors thus attributed the age-related decrease in gyrification in humans as a being related to the extended lifespan of humans, though many other factors also may be relevant (e.g., exercise and diet).

### Non-aging factors that influence brain morphology

While the goal of the current work was to examine the relationship between age and brain morphology, a multitude of other factors are also known to influence estimates of brain morphology. Broadly, these other factors can be categorized as: (1) inter-individual differences; (2) transient physiological changes; and (3) scan-related factors, as outlined in Figure 3. Here we only included age as a predictor and did not include these additional factors, and thus prediction accuracy was inherently limited; however, future work including some of these factors should yield more precise predictions. Furthermore, an important next step will be to examine the relative contribution of different inter-individual difference measures to age-prediction performance.

Inter-individual differences in morphology can arise from many sources, including biological and lifestyle/experiential factors. It is well established that there are sex differences in brain morphology (Barnes et al., 2010; Herron et al., 2015; Hutton et al., 2009; McKay et al., 2014; Potvin et al., 2016; Salat et al., 2004; Sowell et al., 2007). Brain morphology has also been linked with genetic variations; even in healthy individuals, genetic variants of *APOE* (Donix et al., 2010; Honea et al., 2010; Mormino et al., 2014; Okonkwo et al., 2012; Reinvang et al., 2013; Riedel et al., 2016) and *BDNF* (Nemoto et al., 2006; Pezawas et al., 2004; but see Harrisberger et al., 2014) relate to brain morphology estimates. Factors related to perinatal development such as birth weight and nutrition are also associated with morphology, during development and through to adulthood (Strømmen et al., 2015; Walhovd et al., 2012, 2014, 2016). Other lifestyle and experiential factors also can influence brain morphology. Expertise within specific domains has been associated with regional differences in morphology (see May, 2011, for a review), such as in chess players (Hänggi et al., 2014), taxi drivers (Maguire et al., 2000), musicians (Tervaniemi, 2009), athletes (Schlaffke et al., 2014; Tseng et al., 2013), and video game players (Erickson et al., 2010; Kühn et al., 2013). Morphological changes can also arise from experiences such as meditation (Luders et al., 2016; Tang et al., 2015) or from a lack of normal experience, as in acquired blindness (Li et al., in press). Other important inter-individual factors lay somewhere on a continuum in-between biological and lifestyle. Physical health factors, such as exercise and diet, can also influence brain morphology (Booth et al., 2015; Fletcher et al., 2016; Hayes et al., 2014; Kahn et al., 2015; Kullman et al., 2016; Steffener et al., 2016; Williams et al., 2017). Inter-individual differences such as cognitive abilities, personality, and affective style are also associated with morphological differences (Bjørnebekk et al., 2013; Gignac & Bates, 2017; Holmes et al., 2016; Kievit et al., 2014; Riccelli et al., 2017; Valk et al., 2017; Yamagishi et al., 2016). A growing literature has also demonstrated relationships between socioeconomic status and brain morphology (Brito et al., 2017; Brito & Noble, 2014; Hanson et al., 2013; LeWinn et al., 2017; Piccolo et al., 2016).

Beyond inter-individual differences in morphology, transient physiological effects can also influence estimates of brain morphology, and likely also estimates of test-retest reliability. These changes include time-of-day effects (Nakamura et al., 2015; Trefler et al., 2016) and structural changes over the course of longer periods (i.e., infradian), such as the menstrual cycle (fgeern et al., 2011; Lisofsky et al., 2015; Ossewaarde et al., 2013; Protopopescu et al., 2008). Hydration can also influence brain morphometry (Duning et al., 2005; Kempton et al., 2009; Nakamura et al., 2014; Streitbürger et al., 2012; but see Meyers et al., 2016).

Scan-related effects can affect estimates of morphology, without an actual change in morphology. For instance, head movement during scan acquisition can lead to decreased estimates of cortical thickness (Alexander-Bloch et al., 2016; Pardoe et al., 2016; Reuter et al., 2015; Savalia et al., 2017). Effects of head movement on morphology estimates is particularly relevant here, as there is evidence that older adults tend to move more during scanning than younger adults (Andrews-Hanna et al., 2007; Salat, 2014; Savalia et al., 2017; Van Dijk et al., 2012; but see preliminary evidence that fractal dimensionality may be robust to head movement, Madan & Kensinger, 2016). Differences in pulse sequence, magnetic field strength, and other inter-scanner effects can also influence estimates of morphology (Han et al., 2006; Erus et al., 2018; Fortin et al., 2018; Iscan et al., 2015; Jovicich et al., 2009, 2013; Lüsebrink et al., 2013; Madan & Kensinger, 2017b; Potvin et al., 2016; Wonderlick et al., 2009; Zaretskaya et al., 2018), as well as software packages, and even versions, used for data analysis (Chepkoech et al., 2016; Glatard et al., 2015; Gronenschild et al., 2012; Johnson et al., 2017).

In sum, estimates of brain morphology are subject to many sources of variability that are often not considered, with future studies necessary to further our understanding of the complex interplay among these factors.

### Future directions

Prediction accuracy may be improved by extending it to multimodal protocols. For instance, other imaging techniques have been found to index age-related differences, such as iron content (Acosta-Cabronero et al., 2016; Bartzokis et al., 1994; Callaghan et al., 2014; Daugherty & Raz, 2015; Ghadery et al., 2015; Ward et al., 2014; Zecca et al., 2004), white-matter tract integrity (Bender et al., 2016; Chen et al., 2013; Cox et al., 2016; Gunning-Dixon et al., 2009; Kodiweera et al., 2016; Lebel et al., 2012; Teipel et al., 2014), and arterial-spin labeling (Chen et al., 2013). Functional connectivity (e.g., from resting-state fMRI data) could also be used to complement the current approach (e.g., Dosenbach et al., 2010; Liem et al., 2017; Geerligs & Tsvetanov, 2016; Qin et al., 2015). Other non-imaging measures, such as lifestyle factors (e.g., diet, smoking, vascular health, education, and exercise) or blood biomarkers may also improve prediction accuracy (Geerlings & Tsvetanov, 2016; Madsen et al., 2016).

Future work can take advantage of this framework to compare relative differences in ‘brain age’ associated with different participant samples, both healthy and patient populations. For instance, prior studies have found evidence suggesting overestimated aging predictions in patients with Alzheimer’s disease (Franke & Gaser, 2012), traumatic brain injuries (Cole et al., 2015), and schizophrenia (Schnack et al., 2016). Other individuals, such as long-term meditation practitioners, have been found to have significantly underestimated age predictions (Luders et al., 2016).

### Conclusion

Here we demonstrate that reliable age predictions can be made from structural MRI volumes. A combination of cortical thickness and fractal dimensionality yielding the best predictions. The present framework may also prove useful for future research contrasting healthy aging relative to samples of participants exhibiting pathological aging or superagers, as well as for investigating other sources of variability within healthy older adults (such as cognition-, lifestyle-, or health-related factors).

## Acknowledgements

Portions of this research were supported by a grant from the National Institutes of Health (MH080833; to E.A.K.) and by funding provided by Boston College. C.R.M. was supported by a fellowship from the Canadian Institutes of Health Research (FRN-146793).

MRI data used in the preparation of this article were obtained from several sources, data were provided in part by: (1) the Open Access Series of Imaging Studies (OASIS; Marcus et al., 2007); (2) the Information extraction from Images (IXI) dataset (funded by Engineering and Physical Sciences Research Council [EPSRC] of the UK [EPSRC GR/S21533/02]); (3) wave 1 of the the Dallas Lifespan Brain Study (DLBS) led by Dr. Denise Park, and distributed through INDI (Mennes et al., 2013) and NITRC (Kennedy et al., 2016); (4) the BC data were acquired with the support of funding from the Searle Foundation, the McKnight Foundation, and the National Institutes of Mental Health; and (5) cross-sectional aging data acquired by Dr. Craig Stark’s laboratory at University of California–Irvine and distributed through NITRC.

## Author Contributions

C.R.M. conceived the study design, conducted the data analyses, and drafted the manuscript. E.A.K. contributed to the study design and provided critical feedback on the manuscript. Both authors approved of the final version of the manuscript.

## Abbreviations

DKT: Desikan-Killany-Tourville;
FD: fractal dimensionality;
fMRI: functional magnetic resonance imaging;
MdAE: median absolute error;
MRI: magnetic resonance imaging;
PCA: principal component analysis;
RVR: relevance vector regression.

